# Phenotypic and functional characterization of tumor-reactive T cells in malignant pleural effusions

**DOI:** 10.64898/2025.12.05.692662

**Authors:** Noam S. Gaiger, Jack Coburn, Mark N. Lee, Lin Zhang, Heng-Le Chen, Gavitt A. Woodard, Harriet M. Kluger, David A. Schoenfeld, Benjamin Y. Lu

## Abstract

**Background:** Adoptive cell therapy using tumor-infiltrating lymphocytes (TIL) is approved for the treatment of advanced melanoma but is limited by the need for patients to undergo surgical tumor resection. Malignant pleural effusions (MPE) may represent a more accessible source of tumor-reactive T cells. Here, we characterize the cellular composition as well as the transcriptional and functional properties of T cells from MPE compared with pulmonary metastasis and blood from a patient with melanoma.

**Methods:** The immune cellular composition was immunophenotyped by high-dimensional flow cytometry from synchronously collected MPE, a lung metastasis, and blood from a patient with metastatic melanoma. Sorted CD3^+^ T cells were profiled by single-cell RNA sequencing (scRNA-seq) and T cell receptor sequencing (scTCR-seq). TCR reactivity to autologous tumor was evaluated through *in vitro* activation assays with TCR-transduced Jurkat and autologous cancer cells. The killing capacity of *ex vivo* expanded T cells of autologous cancer cells was assessed through *in vitro* cytotoxicity assays.

**Results:** MPE had higher proportions of CD45^+^ immune cells and CD3^+^ T cells (70.5% vs 50%) compared with tumor and was enriched for effector CD8^+^ T cells, CCR7^−^CD45RA^−^ effector memory CD4^+^ T cells, and quiescent CD25^high^CD127^low^ regulatory CD4^+^ T cells. MPE T cells exhibited lower levels of co-inhibitory receptors (PD-1, LAG-3, TIGIT, TIM-3) expression relative to tumor. ScRNA-seq showed enrichment of NK-like effector CD8^+^ T cells in MPE. Pseudotime analysis indicated that MPE T cells were less exhausted than tumor T cells. The clonal repertoire of MPE and tumor highly overlapped, including 62.2% of predicted neoantigen-specific (NeoTCR) clonotypes. Notably, clonally-related NeoTCR T cells in MPE exhibited higher cytotoxic and stemness, and lower exhaustion signatures compared with sister clones in the tumor. Two of four selected NeoTCR clonotypes transduced in Jurkat cells demonstrated MHC class I-restricted reactivity in co-culture with autologous cancer cells. MPE T cells also readily expanded in the presence of high-dose IL-2 and demonstrated MHC class I-dependent killing of autologous cancer cells.

**Conclusions:** MPE harbors polyclonal, tumor-reactive T cells with lower features of terminal exhaustion and higher cytotoxic potential relative to tumor T cells. MPE may therefore serve as a more accessible source for TIL therapy.

## Introduction

Tumor-infiltrating lymphocyte (TIL) therapy leverages the anti-tumor activity of autologous T cells derived from a patient’s own tumor and induces meaningful clinical responses in advanced metastatic melanoma (1–4). A commercially manufactured therapy, lifileucel, is now FDA-approved for the treatment of relapsed advanced cutaneous melanoma (5).

Current methods for generating TIL therapies necessitate surgical tumor resection, which increases the chance of capturing sufficient tumor-reactive T cells but carries procedural risks and delays treatment initiation. Furthermore, in up to 20% of cases, viable TILs cannot be successfully expanded following surgical resection, either due to an insufficient number of pre-existing T cells or diminished proliferation capacity (6). A more accessible source with potentially greater proliferative potential could therefore decrease the time to treatment and improve the accessibility of TIL therapies.

Malignant pleural effusions (MPE) are a complication of advanced cancers and develop when tumors metastasize to the pleural lining. MPE can be sampled through thoracentesis, a common bedside procedure. Prior studies examining the cellular composition of MPE suggest that activated T cells may be present, including a subset of tumor antigen-specific T cells (7, 8). However, the frequency and proliferative potential of these tumor antigen-specific T cells compared to those in the tumor is unknown.

Herein, we systematically characterize the cellular composition and phenotype of synchronously collected MPE, pulmonary metastasis, and blood from a patient with metastatic cutaneous melanoma to define the transcriptional features, clonal repertoire, tumor specificity, and functional reactivity of MPE T cells relative to solid tumor and blood.

## Methods

### Human specimens

Tumor, MPE and blood were collected from a patient with metastatic melanoma under the Yale University Institutional Review Board protocol HIC# 0608001773. Samples were processed using standard density-gradient separation, enzymatic tumor digestion, and cryopreservation protocols. Detailed clinical information, sample processing, experimental, and analytical procedures are provided in the Supplemental Methods.

### Flow cytometry

Samples were stained with viability dye and fluorophore-conjugated antibodies (**Table S4**) and analyzed on a spectral flow cytometer at the Yale Flow Cytometry Core. Live CD45⁺CD3⁺ cells were isolated by spectral cell sorting for single-cell sequencing.

### *Ex vivo* T cell expansion

T cells from MPE and tumor fragments were expanded *ex vivo* in high-dose recombinant human IL-2 under standard culture conditions, as previously described (9).

### Tumor cell line

A patient-derived melanoma cell line was established from surgical tumor tissue and maintained under standard culture conditions.

### TCR Engineering and T-Cell Reactivity Assays

Lentiviral transduction of wild-type B2M and TCRα/β constructs was performed using standard packaging in 293T cells (pLX301, Addgene) (10). Tumor HLA class I expression was assessed by immunofluorescence microscopy, Jurkat T-cell activation was measured by CD69 upregulation following tumor co-culture, and autologous MPE-derived CD8⁺ T-cell reactivity was evaluated by viability and apoptosis after tumor co-culture.

### Single-cell RNA-sequencing and TCR repertoire analysis

Single-cell RNA-seq and TCR libraries were generated using 10x Genomics 5ʹ Gene Expression and V(D)J kits and processed with CellRanger (GRCh38). Downstream analysis, including QC, normalization, batch correction, clustering, differential expression, pseudotime, pathway enrichment, and TCR analysis, was performed using the open-source Scanpy and scVI-tools, Palantir, GSEApy, and Scirpy packages. Tumor-reactive T cells were identified by AUCell enrichment of NeoTCR signatures (11–16).

### Statistical analysis

All statistical analyses were performed using GraphPad Prism (v10.5.0) and Python (v3.10.16). Flow cytometry data were compared across groups using one-way ANOVA with Tukey’s multiple comparisons test. Differences in gene signature scores between compartments were assessed using pairwise Mann–Whitney U tests. All *p*-values are two-sided, with statistical significance defined as p < 0.05.

## Results

### MPE is enriched with activated, effector T cells which exist in a pro-inflammatory immune microenvironment

To characterize the cellular composition of immune cells across tissue compartments, we used spectral flow cytometry to analyze the expression of sixteen cell lineage markers and sixteen T cell markers in MPE, lung metastasis (“Tumor”), and blood (**Figure 1A; Figure S1-S4**). Overall, MPE had the highest percentage of CD45^+^ immune cells (94.8% in MPE vs. 89.8% in blood vs. 14.7% in tumor; **Figure S5**) and CD3^+^ T cells (74.9% in MPE vs. 58.6% in tumor vs. 48.7% in blood; **Figure 1B**). Of CD3^+^ T cells, the tumor had the highest proportion of CD8^+^ T cells (55.8% in tumor vs. 29.9% in MPE vs. 31.6% in blood), which were predominantly composed of CD45RA^−^CCR7^−^ effector memory and CD45RA^+^CCR7^−^ terminally differentiated (Temra) subsets (**Figure 1C**). Tumor CD8^+^ T cells were also most enriched with CD103^+^CD69^+^ resident memory T cells (Trm; 24.6% in tumor vs. 8.4% in MPE vs. 0.3% in blood; **Figure 1D**) and exhibited the highest expression of the co-inhibitory receptors PD-1, LAG-3, TIGIT, and TIM-3 (**Figure 1H**; **Figure S6A**). In contrast, MPE CD8^+^ T cells had the highest proportion of CX3CR1^+^KLRG1^+^ effector T cells (5.6% in MPE vs. 2.9% in tumor vs. 2.8% in blood), the lowest proportion of CD45RA^+^CCR7^+^ naïve T cells (Tn; 7.9% in MPE vs. 9.6% in tumor vs. 22.7% in blood), and intermediate expression of multiple co-inhibitory receptors (**Figure 1C, 1E, 1H; Figure S6A**).

**Figure 1.**
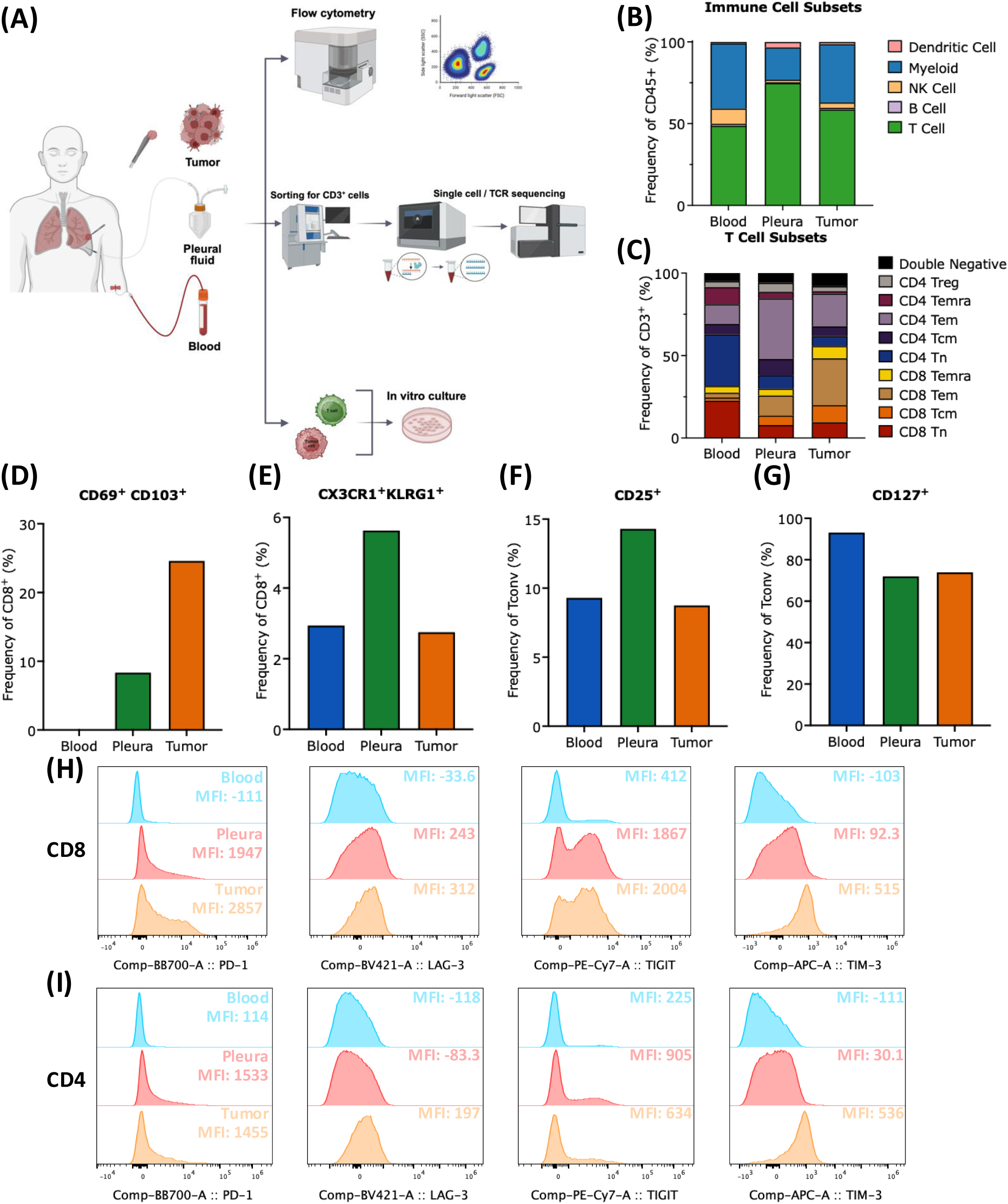
Immunophenotyping of T cells across tissue compartments. **(A)** Design and analytical pipeline of multiparametric flow cytometer, 5’ droplet-based scRNA-seq with paired TCR sequencing, and functional co-culture assay to characterize T cells in blood, malignant pleural effusion (MPE), and a pleural metastasis (tumor). **(B)** Proportion of dendritic cells, monocytes, NK cells, B cells, and T cells across tissue compartments as a percentage of *CD45^+^* cells. **(C)** Proportion of *CD8^+^* or *CD4^+^ CD45RA^+^CCR7^+^* naïve (Tn), *CD45RA^-^CCR7^+^* central memory (Tcm), *CD45RA^-^CCR7^-^* effector memory (Tem), *CD45RA^+^CCR7^-^* terminally differentiated (Temra), and *CD4^+^CD25^high^CD127^low^* regulatory (Treg) T cells across tissue compartments as a percentage of *CD3+* cells. **(D)** Frequency of *CD8*^+^*CDcS^+^CD103^+^* T cells across tissue compartments. **(E)** Frequency of *CD8*^+^*CX3CR1^+^KLRG1^+^* T cells across tissue compartments. **(F)** Frequency of *CD25*^+^ and **(G)** *CD127*^+^ on conventional (Tconv) T cells. **(H)** Histograms and mean fluorescence intensity of *PD-1*, *LAG-3*, *TIGIT*, and *TIM-3* on *CD8*^+^ T cells and **(I)** Tconv across tissue compartments.

Despite a similar proportion of CD4^+^ T cells between MPE and blood, MPE CD4^+^ conventional T cells (Tconv) displayed a more activated and effector phenotype. CD4^+^ Tconv in the blood contained the highest proportion of Tn (31.2% in blood vs. 8.2% in MPE vs. 5.9% in tumor) and the highest expression of the memory marker CD127 (93.1% in blood vs. 72.0% in MPE vs. 73.9% in tumor; **Figure 1C, 1G**). In contrast, Tconv cells in MPE contained the highest proportion of Tem cells (72% in MPE vs. 63% in tumor vs. 20.1% in blood), the highest expression of the activation marker CD25 (14.3% in MPE vs. 9.3% in blood vs. 8.8% in tumor), intermediate expression of the activation marker CD69 (5.61% in MPE vs. 0.5% in blood vs. 57.8% in tumor), and intermediate expression of PD-1, LAG-3, and TIM-3, which is overall consistent with an activated, effector phenotype (**Figure 1C**, **1F**, **1I**; **Figure S6B**).

While the proportion of CD4^+^CD25^high^CD127^low^ regulatory T cells (Treg) was numerically highest in MPE (6.9% in MPE vs. 3.8% in tumor vs. 4.2% in blood), MPE Tregs possessed lower levels of activation markers (CD69, HLA-DR) and co-inhibitory receptor (PD-1, LAG-3, TIM-3) expression (**Figure 1C**, **Figure S5J-M**, **Figure S7A**). Taken together, these findings suggest that Tregs in MPE have a less activated and less suppressive phenotype than tumor Tregs.

Among non-T cell populations, MPE was enriched for CD14^+^CD16^+^ monocytes (59.3% in MPE vs. 27.4% in blood vs. 1.9% in tumor; **Figure 1B**, **Figure S5B**). In addition, NK cells within MPE were enriched for CD56^high^CD16^-^ NK cells (20.2% in MPE vs. 2.0% in blood vs. 0.5% in tumor; **Figure S5C**) which are associated with effector cytokine production (17). These results suggest that MPE possesses a unique immunological milieu with activated effector CD8^+^ and CD4^+^ T cells that are less exhausted than tumor T cells.

### MPE T cells exist in an intermediate transcriptional state between blood and tumor T cells

To define transcriptional features of T cells between tissue compartments, we performed scRNAseq paired with scTCRseq on sorted CD45^+^CD3^+^ T cells. In total, we analyzed 9,819 high-quality T cells, including 3,156 from MPE, 3,553 from tumor, and 3,110 from blood. Overall, thirteen distinct transcriptomic clusters were identified, including seven CD8⁺ T cell clusters, five CD4⁺ T cell clusters, and one proliferating cluster (**Figure 2A-B, Figure S8, Table S1-2**).

**Figure 2.**
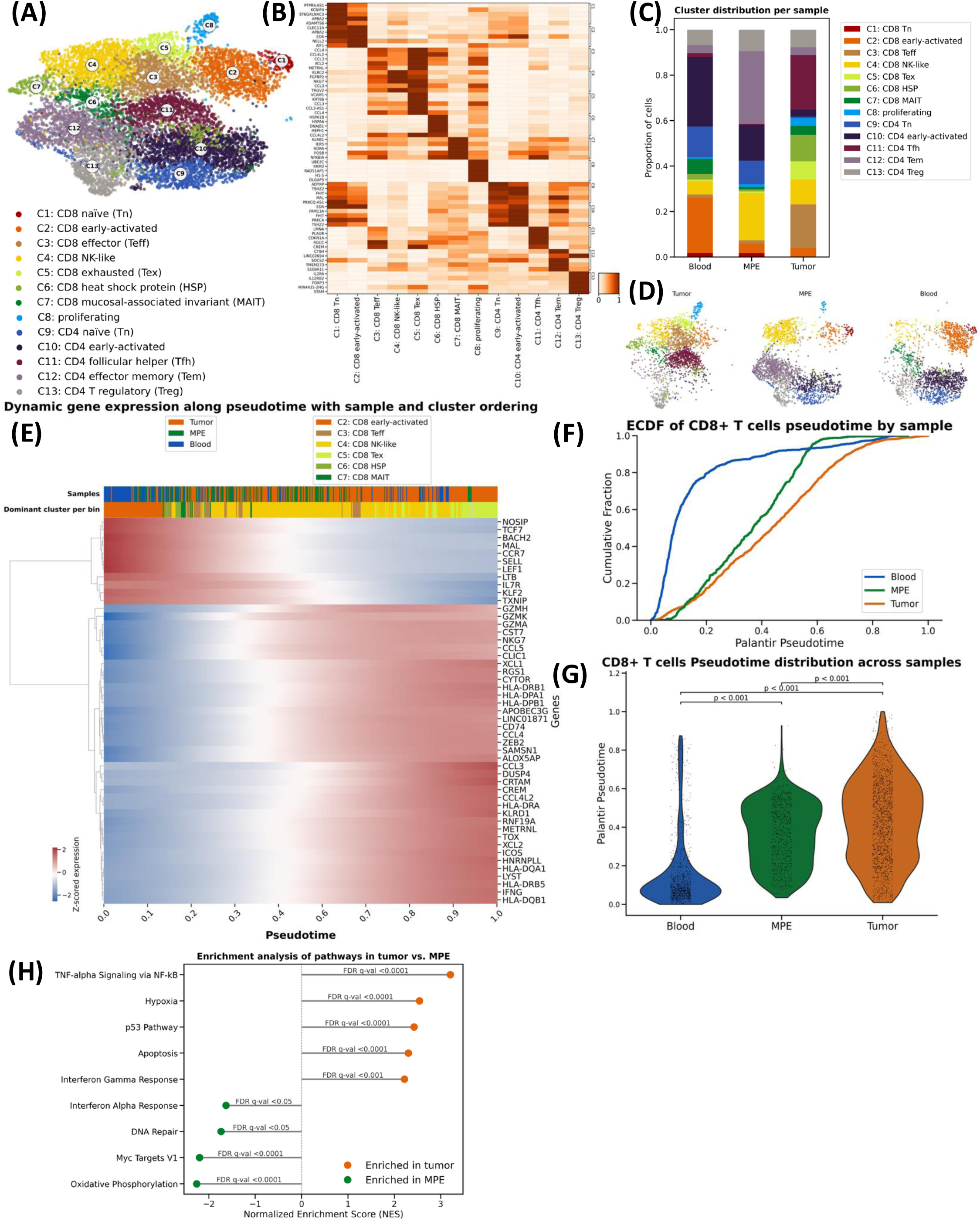
Transcriptional profiles and dynamics of T cells across tissues. **(A)** UMAP visualization of single-cell transcriptomic data from 9,819 T cells isolated from tumor, malignant pleural effusion and peripheral blood. Clusters were annotated using published scRNA-seq gene signatures (Table S2). (**B**) Expression matrixplot of differentially expressed genes across cell clusters **(C)** Distribution of clusters across samples **(D)** Compartment-specific UMAP visualization of T cell clusters **(E)** Differential gene expression along pseudotime trajectory **(F)** Empirical cumulative distribution function (ECDF) of compartment-specific CD8⁺ T cells along the pseudotime trajectory. **(G)** Violin plot of cells per-sample scattered along pseudotime. **(H)** Gene set enrichment analysis (MSigDB Hallmark 2020) identifies pathway differences between tumor and MPE samples.

Compositional analysis of transcriptional cluster distributions between tissues were overall consistent with the flow cytometry findings: naïve and early activated CD8⁺ and CD4⁺ T cell populations (C1, C2, C9, C10) were most enriched in the blood (**Figure 2C-D, Table S3**); CD8⁺ NK-like (C5) T cells and CD4⁺ Tem (C12) cells were enriched in MPE (C5: 20.3% of MPE T cells, C12: 33.6% of MPE); and CD8⁺ effector (C3), CD8⁺ exhausted (C4) T cells, and Tfh (C11) were enriched in tumor (C3: 19.6% of tumor T cells, C4: 7.3% of tumor T cells, C11: 26.7% of tumor T cells). Proliferating T cells (C8) were rare in blood (0.2%) and most abundant in tumor (3.4%) (**Figure 2C-D, Table S3**).

To understand the relative transcriptional states along a continuous differentiation trajectory, we next performed pseudotime trajectory analyses of CD8^+^ and CD4^+^ T cells separately (**Figure 2E, Figure S9**). Dynamic gene expression analysis of CD8^+^ T cells demonstrated a continuum between genes associated with naïve states (*BACH2, CCR7, SELL, TCF7*) to genes associated with highly activated states (*GZMK, NKG7, CCL5, HLA-DPA1, HLA-DRB1*) (**Figure 2E**). Notably, unsupervised cell ordering showed that T cells segregated by tissue of origin across the pseudotime trajectory. Blood T cells predominantly occupied earlier pseudotime values relative to MPE and tumor (*p* < 0.001; **Figure 2F-G**). In contrast, tumor T cells were positioned at the terminal end of the dynamic trajectory, with significantly higher pseudotime values than both MPE (*p* < 0.001) and blood (*p* < 0.0001; **Figure 2F-G**). Gene set enrichment analysis revealed that both MPE and tumor were enriched for inflammatory pathways indicative of T cell activation (**Figure 2H**). However, tumor-derived T cells were also enriched for chronic stimulation and exhaustion pathways, including apoptosis and hypoxia (**Figure 2H**). Overall, these results demonstrate that MPE is enriched for activated, effector T cells that have lower transcriptional features of functional exhaustion relative to tumor.

### Predicted tumor-reactive T cells are shared between tumor and MPE

To understand the clonotypic relationship across compartments, we performed scTCRseq and identified 5,717 unique TCR clonotypes. T cells in the tumor exhibited the highest clonality (**Figure 3A-D, Figure S10**), with expanded clonotypes (n ≥ 2) comprising 36.8% of tumor T cells, 22.2% of MPE T cells, and 5.3% of blood T cells (**Figure 3C**). Among 196 expanded clonotypes identified in the tumor, 90 clonotypes (45.9%) were also detected in MPE, whereas only 28 clonotypes (14.3%) were detected in blood. The overall clonal repertoire highly overlapped between tumor and MPE (**Figure 3E-G, Figure S11**). Furthermore, clone sizes in MPE were strongly correlated with those in tumor (**Figure 3F**). Taken together, these data indicate that the MPE T cell repertoire largely reflects the clonal repertoire in tumor.

**Figure 3.**
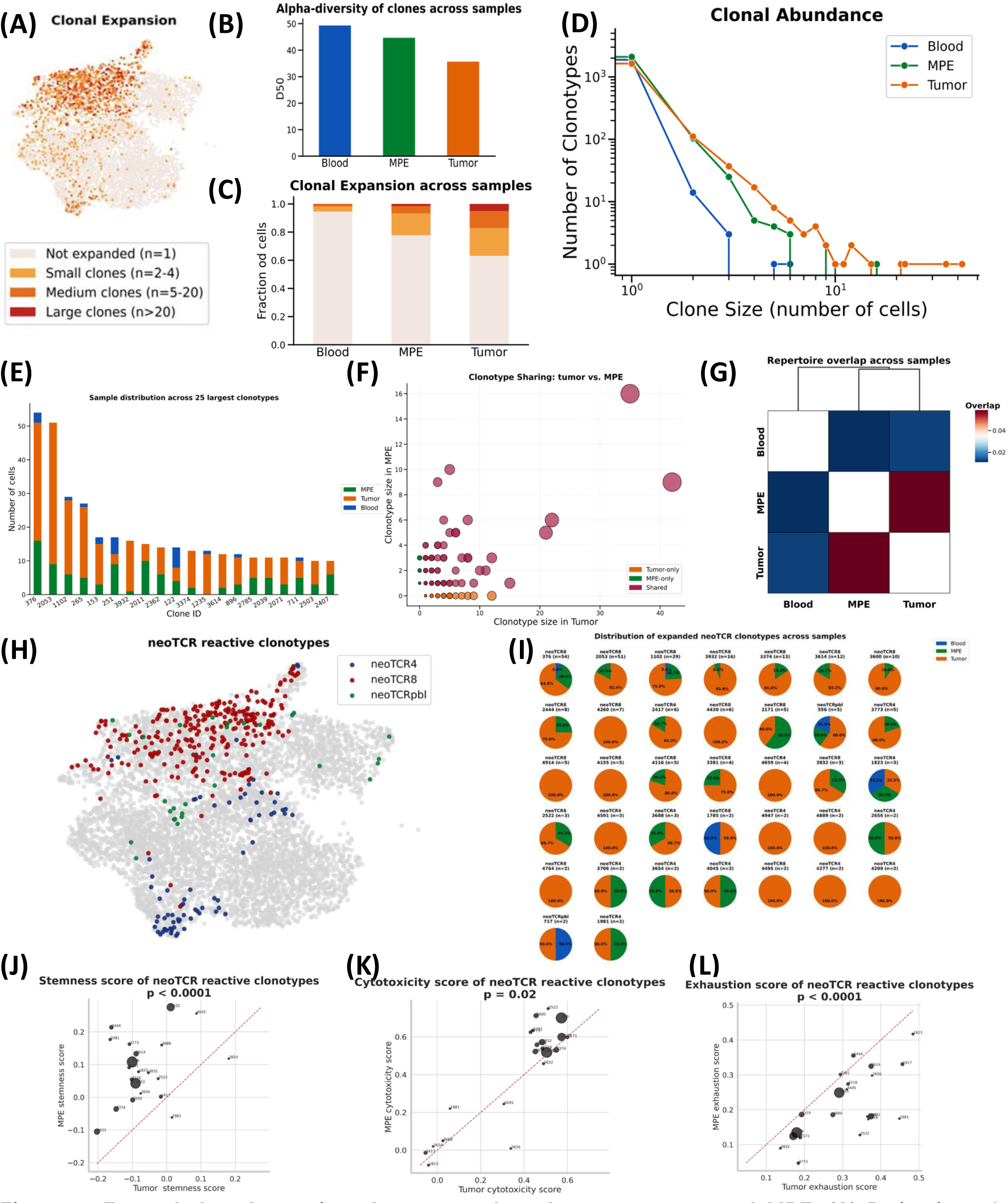
Expanded and reactive clones are shared across tumor and MPE. **(A)** Projection of expanded clonotypes onto the T cell transcriptomic UMAP, categorized by clone size **(B)** Alpha diversity, representing the minimum number of unique clonotypes that account for 50% of all cells **(C)** Clonal expansion across blood, MPE, and tumor **(D)** Distribution of clonotypes by clone size across blood, MPE, and tumor **(E)** Distribution of the top 20 tumor-enriched clonotypes across compartments **(F)** Clonotype sharing between tumor and MPE **(G)** Heatmap of clonotype overlap between blood, MPE, and tumor **(H)** Projection of neoantigen-reactive TCR clonotypes onto the UMAP. NeoTCRs were identified using published gene signatures (neoTCR4, neoTCR8, and neoTCRpbl) (I) Distribution of the top 30 neoTCR clonotypes across compartments. (J–L) Gene signature scores for stemness **(J)**, cytotoxicity **(K)**, and exhaustion **(L)**.

To identify predicted neoantigen-specific T cells, we applied previously described, experimentally-validated gene signatures to the tumor (NeoTCR8 and NeoTCR4) (18) and blood (NeoPBL) (19). This approach identified 118 NeoTCR clonotypes across the three neoantigen gene signatures (collectively referred to as NeoTCR), including 37 (31.4%) expanded clonotypes (n ≥ 2). Tracing TCRs across tissue compartments, we were then able to identify clonally-related NeoTCR T cells in the MPE. 23 of the 37 expanded NeoTCR clonotypes in tumor (62.2%) were also present in MPE (**Figure 3I**). In contrast, among the 37 expanded NeoTCR clonotypes, only 6 (16.2%) were observed in blood (**Figure 3I**). Overall, 14.9% and 6.0% of tumor and MPE CD8^+^ T cells, respectively, were predicted to be tumor reactive, compared to only 0.6% of CD8⁺ T cells in the blood.

Next, we compared the transcriptional states of each NeoTCR shared between tumor and MPE using previously described stemness, cytotoxicity and exhaustion signatures (**Figure 3J-3L**) (20). NeoTCR clonotypes in MPE had higher stemness (*p* < 0.0001) and cytotoxicity scores (*p =* 0.02), while clonotypes in the tumor had significantly higher exhaustion scores (*p* < 0.0001). Notably, NeoTCR clonotypes in MPE also had a higher expression of CD39⁻CD69⁻ stem-like signature (*p* < 0.05), which has been associated with durable responses with TIL therapy. In contrast, NeoTCR clonotypes in tumor more highly expressed a CD39⁺CD69⁺ terminal differentiation signature associated with poor responses (p < 0.001; **Figure S12**). While these gene signatures were originally defined in expanded TILs, these findings demonstrate that predicted tumor-specific T cells in MPE are in a less terminally exhausted differentiation state than tumor and support the therapeutic potential of MPE-derived T cells for ACT.

### Shared NeoTCR8 clonotypes show functional reactivity to autologous cancer cells

To functionally confirm the tumor reactivity of NeoTCR8 clonotypes, we first established an autologous cancer cell line from the lung metastasis (**Figure 4A**). We noted a paucity of HLA class I expression on the cancer cell line (**Figure 4B**) and identified a single-nucleotide variant resulting in a missense mutation in the *B2M* gene (c.92C>G) (21) on whole-exome sequencing analysis. HLA class I expression was restored by transducing the *B2M* gene but not *HLA-A2* (**Figure 4B**). We next selected four TCR’s that were predicted to be tumor reactive based on NeoTCR8 expression, had the highest degree of clonal expansion, and were detected in both tumor and MPE (**Figure 4C-D**). Paired TCRα and TCRβ sequences of each selected clonotype were then cloned into CD8^+^ TCR knockout Jurkat T cells, as previously described (10), and co-cultured with an autologous cancer cell line. Two of the four selected TCRs (TCR1, TCR4) resulted in significant increases in the activation marker CD69 when specifically co-cultured with B2M-transduced, HLA class I-expressing cells but not control or HLA-A2 transduced cancer cells (**Figure 4E-F, Figure S13**).

**FIGURE 4.**
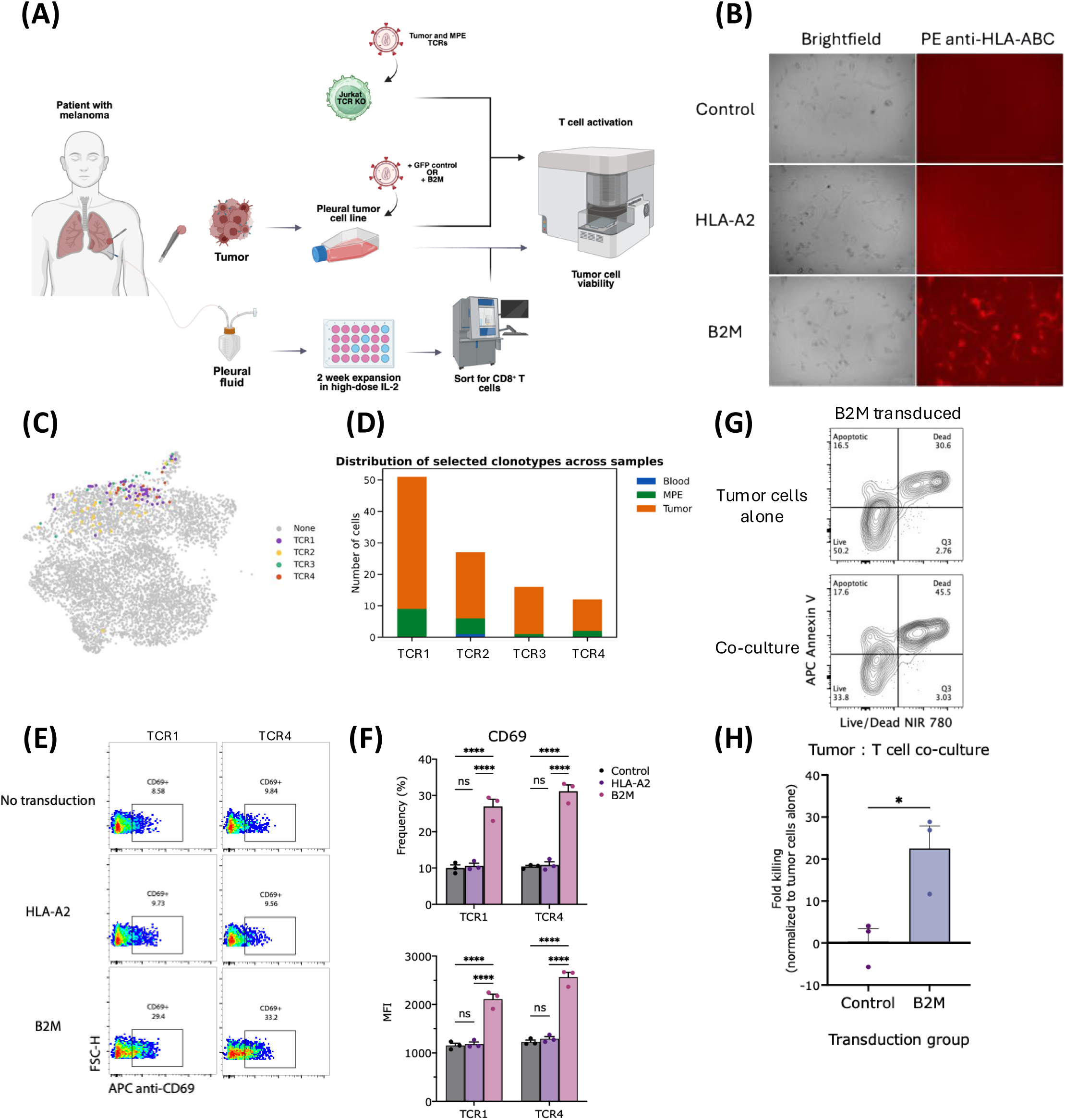
Functional specificity and reactivity of MPE T cells. **(A)** Diagram of ex vivo tumor and TIL expansion and co-culture assay. **(B)** Brightfield and immunofluorescence image of cancer cells with or without B2M or HLA-A2 transduction. **(C)** Dimensionality reduction UMAP plot highlighting selected CD8+ T cells frequency. **(D)** Stacked bar plot of TCR abundance and across which tissues. Representative **(E)** flow cytometry plots or **(F)** summary plots showing CD69 expression. Representative **(G)** flow cytometry plot and **(H)** summary bar plot of T cell-directed tumor cell killing. Between-group differences are measured by ordinary one-way ANOVA with Tukey’s multiple comparisons test (*p < 0.05, **p < 0.01).

### *Ex vivo* expanded MPE T cells exhibit cytotoxic capacity against autologous cancer cells

To assess the cytotoxic potential of *ex vivo* expanded MPE T cells, we next cultured unselected cells from MPE in high-dose IL-2 (HD IL-2) for up to two weeks, similar to conventional TIL expansion methods (**Figure 4A**, **Figure S14**) (9). Notably, tumor-infiltrating T cells failed to expand from tumor chunks cultured in a similar manner, possibly due to their terminal exhaustion state or a lack of sustained TCR signaling in the presence of class I deficient cancer cells. To examine the cytotoxic capacity of expanded MPE T cells, we co-cultured sorted CD8^+^ T cells from expanded MPE cells with CFSE-labeled, autologous cancer cells transduced with GFP or B2M. Following 20 hours in co-culture, we observed significantly higher cancer cell killing when T cells were co-cultured with B2M transduced tumor cells (*p* = 0.035; **Figure 4G-H**). These data demonstrate the feasibility and cytotoxic capacity of collecting and expanding tumor reactive T cells from MPE.

## Discussion

The polyclonal nature and enrichment for tumor antigen-specific T cells within the tumor microenvironment is thought to underly the success of TIL therapy (22). However, chronically stimulated T cells undergo progressive dysfunction – termed T cell exhaustion – and lose effector and proliferative capacity (20, 23, 24). An optimal biological source for TIL therapy therefore balances both tumor antigen-specificity and proliferative capacity.

In this study, MPE was highly enriched for effector T cells with lower exhaustion features compared with tumor T cells and shared highly overlapping clonal repertoire with tumor T cells, including predicted tumor reactive clonotypes. Furthermore, predicted tumor-reactive clonotypes in MPE were enriched for stem-like phenotype associated with durable responses to TIL therapy, whereas predicted tumor-reactive clonotypes in tumor were dominated by a terminally differentiated phenotype associated with poorer outcomes (24).

A major limitation of current TIL therapy is the need for invasive surgical tumor resection, excluding patients who are not surgical candidates. In contrast, MPE can be collected through in-office thoracenteses and may provide an alternative biological source of TIL therapy for patients who develop MPE.

While our findings are encouraging, validation in larger cohorts and across additional cancer types is essential to confirm their generalizability and functional effectiveness of a TIL product derived from MPE *in vivo*. Furthermore, we found that anti-tumor T cells exist in the MPE in an activated transcriptional state, and by tracing clonotypes shared with tumor cells, we infer that they likely recognize the same antigens. However, we cannot exclude the possibility that additional unrecognized anti-tumor T cells exist in alternative phenotypic states within the MPE. Additionally, while our functional assays demonstrated tumor-killing capacity, further studies are needed to validate the functional effectiveness of a TIL product derived from MPE *in vivo*.

In summary, our study demonstrates that MPE harbors tumor-reactive T cells with high cytotoxic tumor-killing capacity and a lower degree of terminal exhaustion relative to tumor. These findings therefore support ongoing efforts to use MPE as a more accessible and novel immunotherapy for patients with pleural metastases (25).

## Declarations

### Ethics approval and consent to participate

This study was approved by the Yale University Institutional Review Board (HIC# 0608001773). Written informed consent was obtained from the patient prior to sample collection.

## Author’s contributions

BYL and DAS conceptualized and supervised the study. GAW acquired tissue samples. LZ, NSG, JC, and BYL processed the samples. JC, BYL, and MNL performed experimental data acquisition. NSG, HLC, and BYL performed computational and statistical analyses. BYL, DAS, JC, NSG, MNL, and HMK interpreted the results. NSG, JC, BYL, and DAS drafted the manuscript. All authors reviewed and approved the final manuscript.

## Availability of data and material

The processed single-cell data, including the annotated h5ad object used for all analyses, have been deposited in Figshare and are available through https://figshare.com/s/4f4090dcc3bea285ad0b.

## Acknowledgments

The authors are grateful to the patient and their family for the donation of tissue samples for the studies presented in this report. The authors would also like to thank the Yale Flow Cytometry Core for their assistance with cell sorting and instrument maintenance, the Keck Microarray Shared Resource (KMSR) at Yale University for their assistance with single-cell library preparation, and the Yale Center for Genome Analysis (YCGA) for next-generation sequencing services.

## Funding sources

This research was supported in part by a Career Enhancement Program Grants from the Yale SPORE in Lung Cancer P50CA196530 (MNL, BYL, GAW), the National Cancer Institute (NCI) Award Number K08CA270191 (MNL), the NCI Award Number K12CA215110 (HMK, BYL), the U.S. Department of Defense KC230193/HT9425-24-1-0759 (DAS), and the Conquer Cancer Career Development Award (BYL). Any opinions, findings, and conclusions expressed in this material are those of the author(s) and do not necessarily reflect those of the American Society of Clinical Oncology® or Conquer Cancer®. The Yale Flow Core is supported in part by an NCI Cancer Center Support Grant National Institutes of Health P30 CA016359. The YCGA is supported by the National Institute of General Medical Sciences of the National Institutes of Health under Award Number 1S10OD030363-01A1.

## Declaration of interest

DAS has received institutional research support from Pfizer and HiberCell, and research materials from Lassen Therapeutics. HMK reports institutional research grants from Pfizer, Merck, Bristol Myers Squibb and Apexigen and consultation fees from Iovance, Celldex, Merck, Elevate Bio, Instil Bio, Bristol Myers Squibb, Clinigen, Shionogi, Chemocentryx, Calithera, Signatero, Gigagen, GI reviewers, Seranova, Pliant Therapeutics and Teva Pharmaceuticals. GAW reports advisory board participation for AstraZeneca and Bristol Myers Squibb. The remaining authors have declared no conflict of interest.

## List of abbreviations

(TIL): Tumor-infiltrating lymphocytes
(MPE): Malignant pleural effusions
(scRNA-seq): Single-cell RNA sequencing
(scTCR-seq): T cell receptor sequencing

