## Supplemental Figure for "Phenotypic and functional characterization of tumor-reactive T cells in malignant pleural effusions"

### Figure S1

**(A)** *PBMCS*

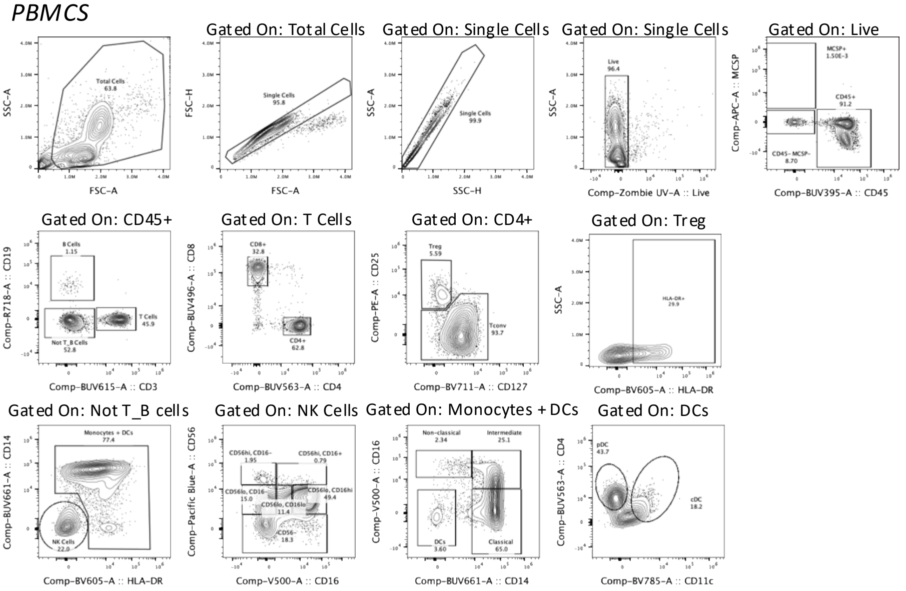


**(B)** *Ex vivo MPE*

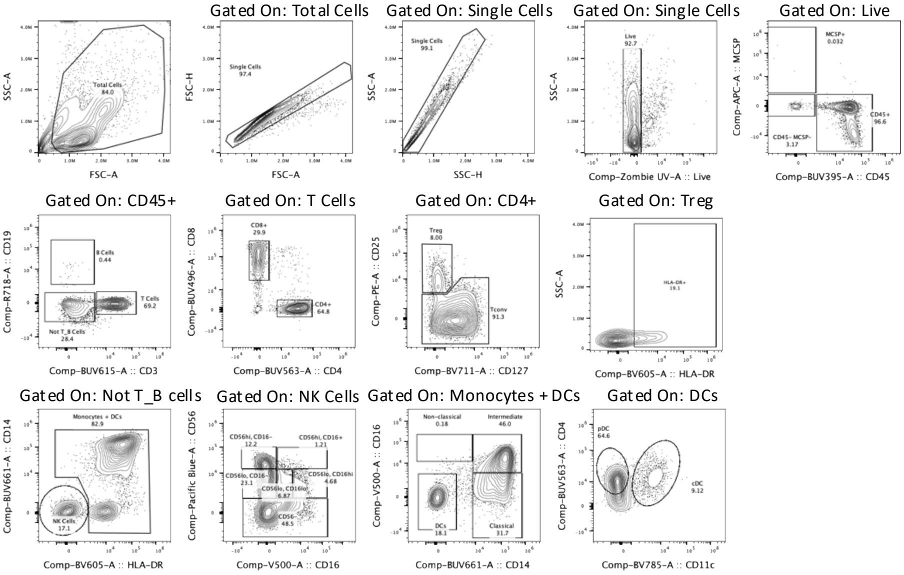


**(C)** *Tumor*

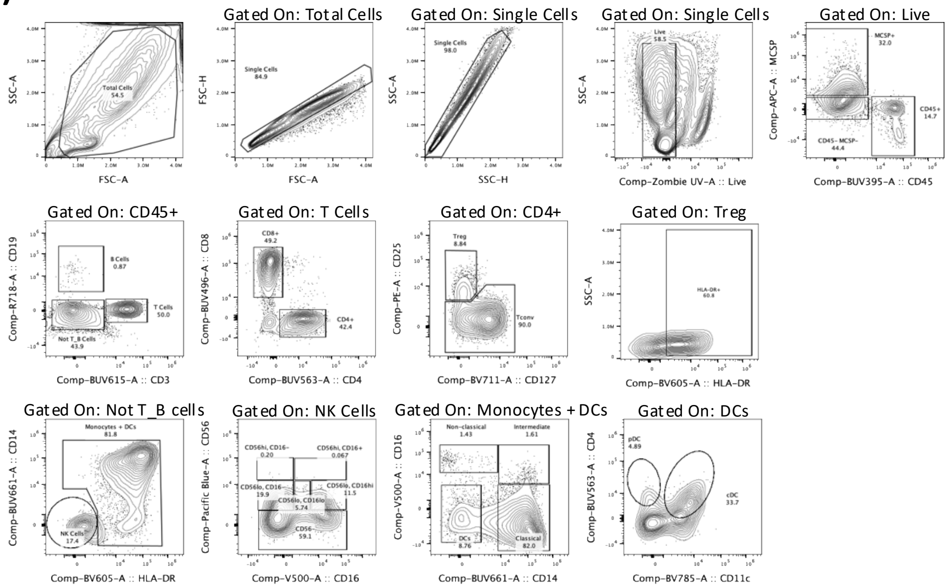


Flow cytometry gating strategy to analyze *ex vivo* immune cell populations in (A) peripheral blood mononuclear cells (PBMC), (B) malignant pleural effusions (MPE), and (C) tumor.

### Figure S2

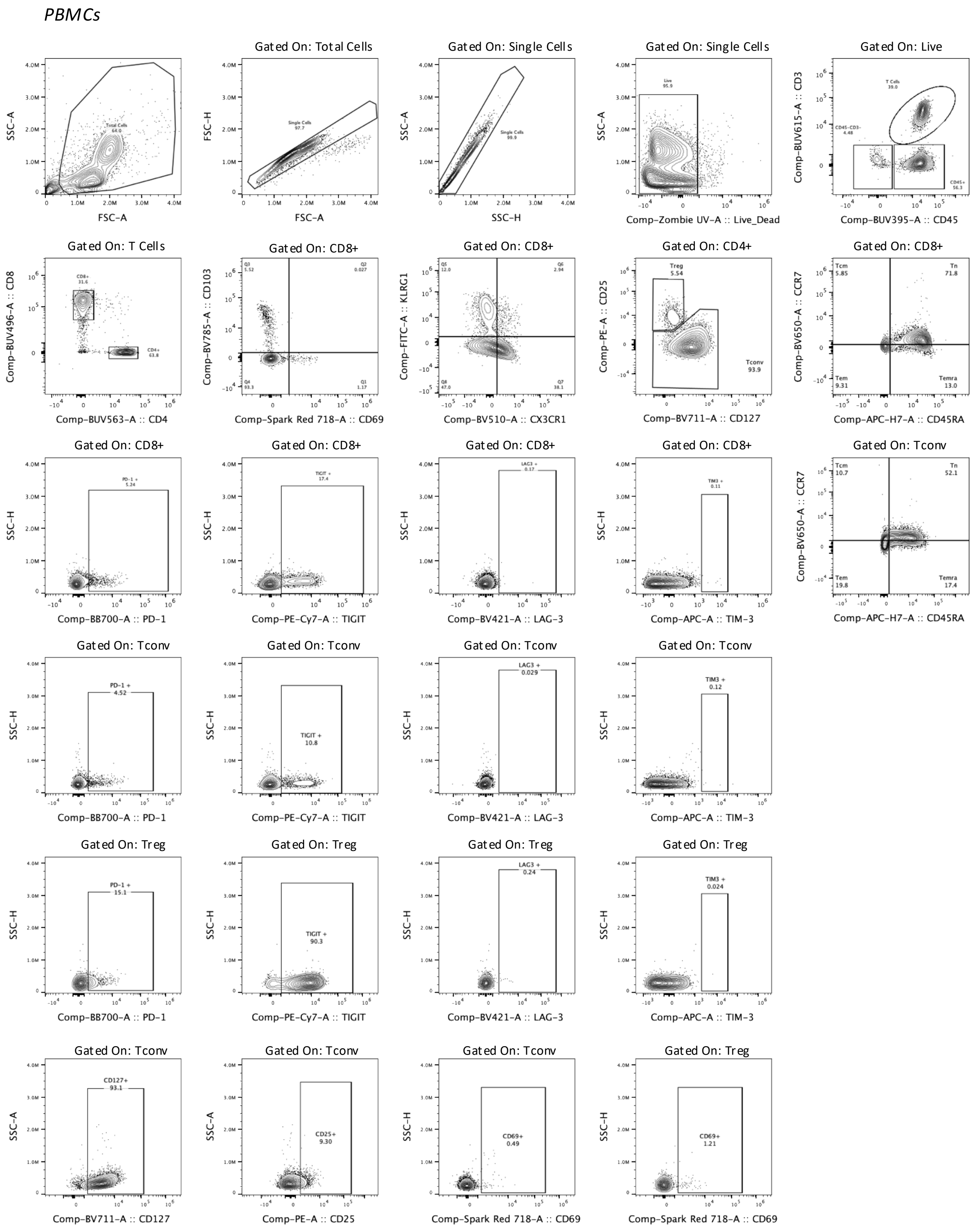


Flow cytometry gating strategy to analyze T cell populations and co-inhibitory receptor expression in PBMC.

### Figure S3

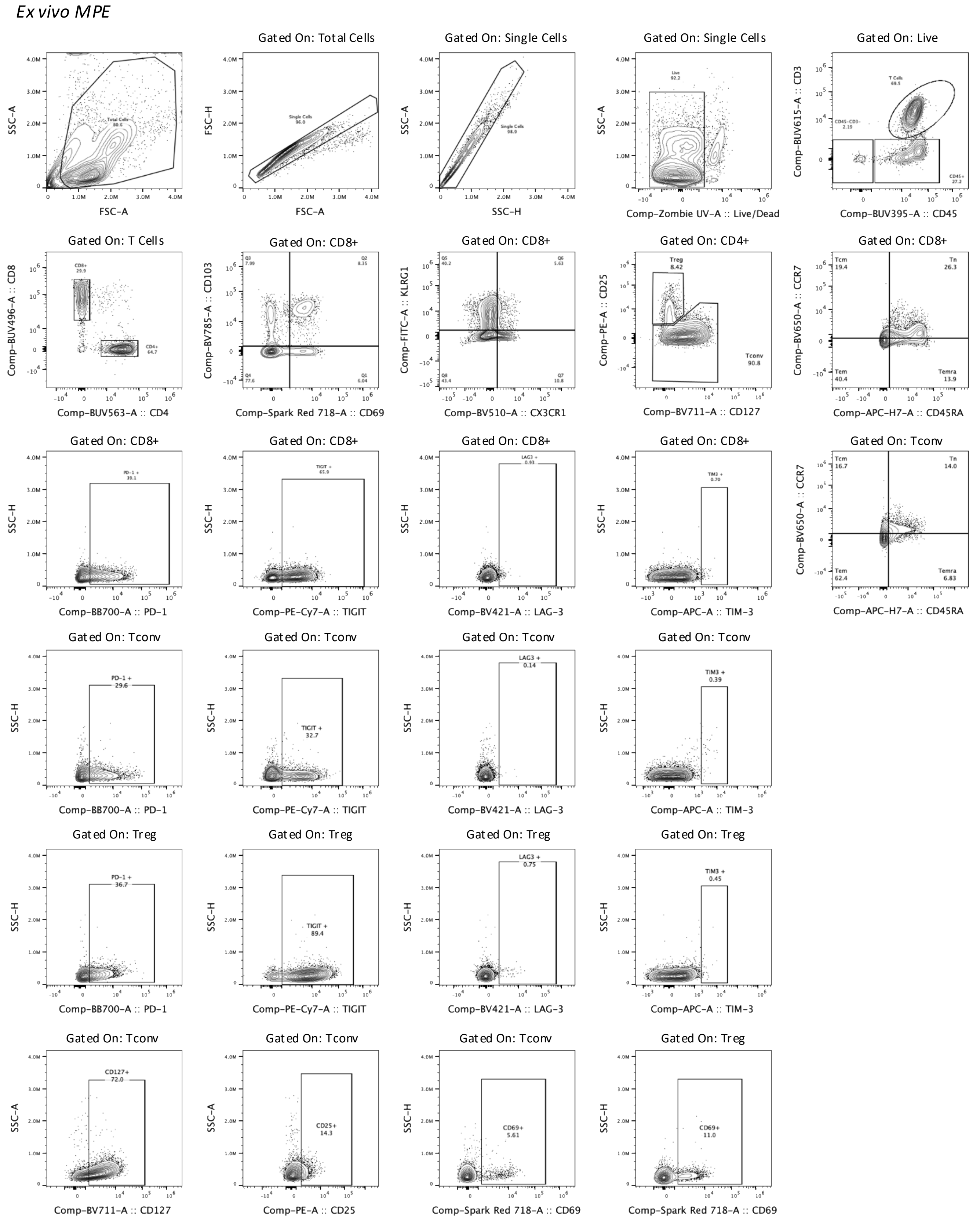


Flow cytometry gating strategy to analyze T cell populations and co-inhibitory receptor expression in *ex vivo* MPE.

### Figure S4

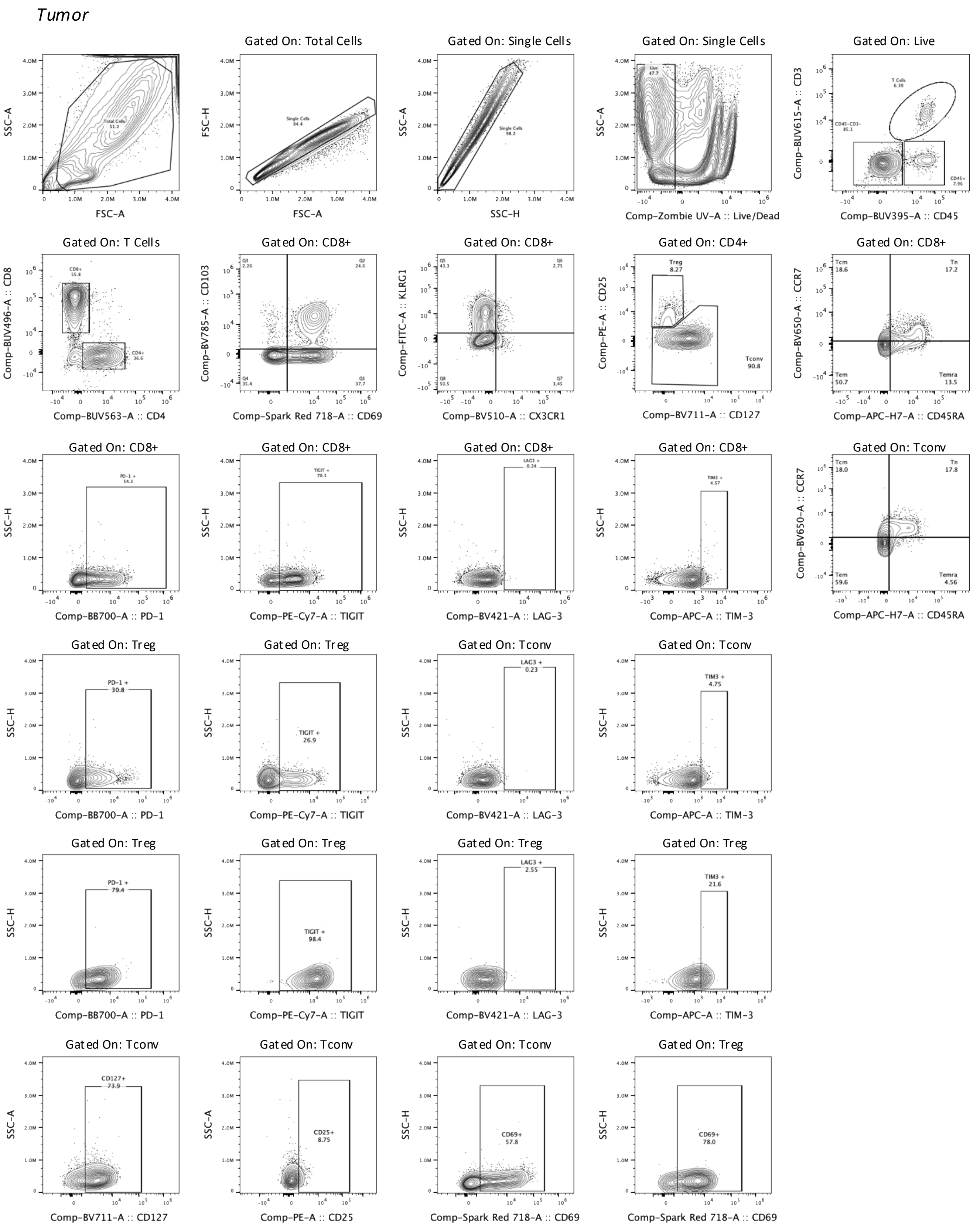


Flow cytometry gating strategy to analyze T cell populations and co-inhibitory receptor expression in tumor.

### Figure S5

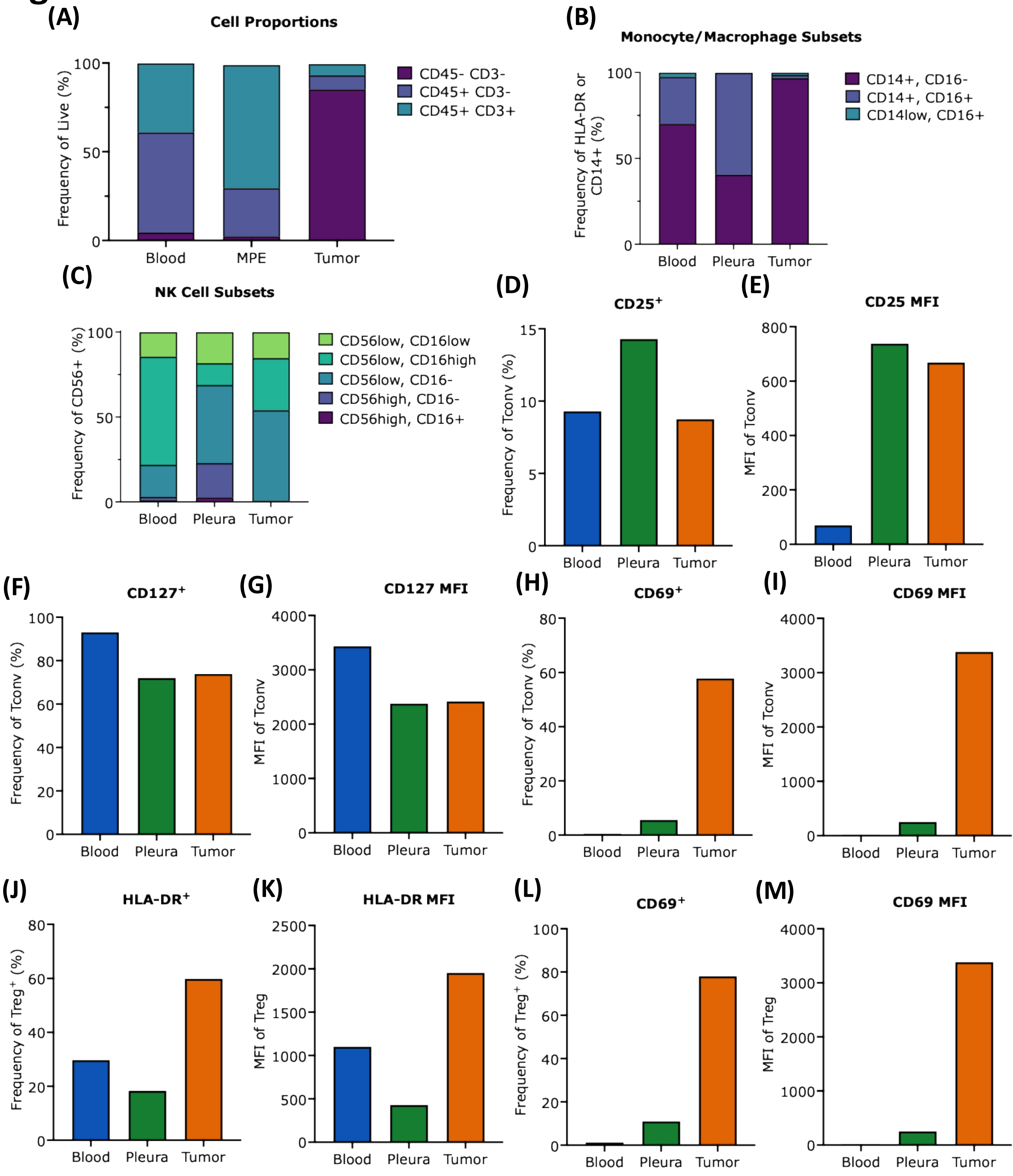


**(A)** Proportion of *CD45$^-$CD3$^-$*, *CD45$^+$CD3$^-$*, and *CD45$^+$CD3$^+$* cells of total live cells across tissue compartments. **(B)** Proportion of *CD14$^+$CD16$^-$*, *CD14$^+$CD16$^+$*, and *CD14$^{low}$CD16$^+$* monocytes/macrophages across tissue compartments as a percentage of total monocytes/macrophages. **(C)** Proportion of *CD56$^{high}$CD16$^-$*, *CD56$^{high}$CD16$^+$*, *CD56$^{low}$CD16$^-$*, *CD56$^{low}$CD16$^{low}$*, and *CD56$^{low}$CD16$^{high}$* NK cell populations across tissue compartments as a percentage of *CD56$^+$* cells. **(D)** *CD25* expression on conventional *CD4$^+$CD25$^{low/–}$*(Tconv) T cells by frequency and **(E)** mean fluorescence intensity (MFI). **(F)** *CD127* expression on Tconv cells by frequency and **(G)** MFI. **(H)** *CD69* expression on Tconv cells by frequency and **(I)** MFI. **(J)** *HLA-DR* expression on regulatory *CD4$^+$CD25$^{high}$CD127$^{low}$* (Treg) T cells by frequency and **(K)** MFI. **(L)** *CD69* expression on Treg cells by frequency and (M) MFI.

### Figure S6

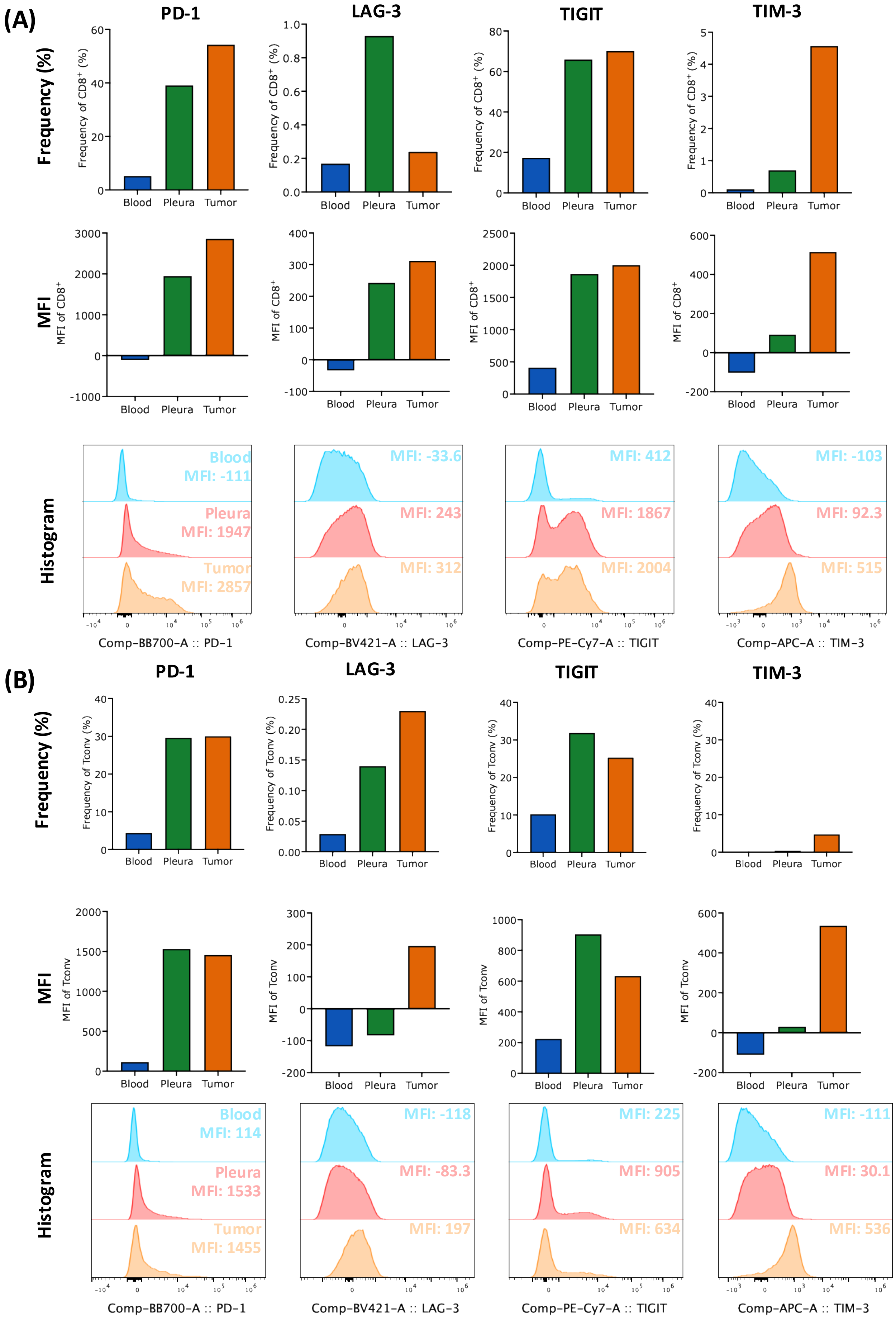


**(A)** Frequency, mean fluorescence intensity (MFI), and histograms showing expression of *PD-1*, *LAG-3, TIGIT*, and *TIM-3* on *CD8⁺* T cells and **(B)** Conventional $CD4^{+}CD25^{low/-}$ (Tconv) T cells across tissue compartments.

### Figure S7

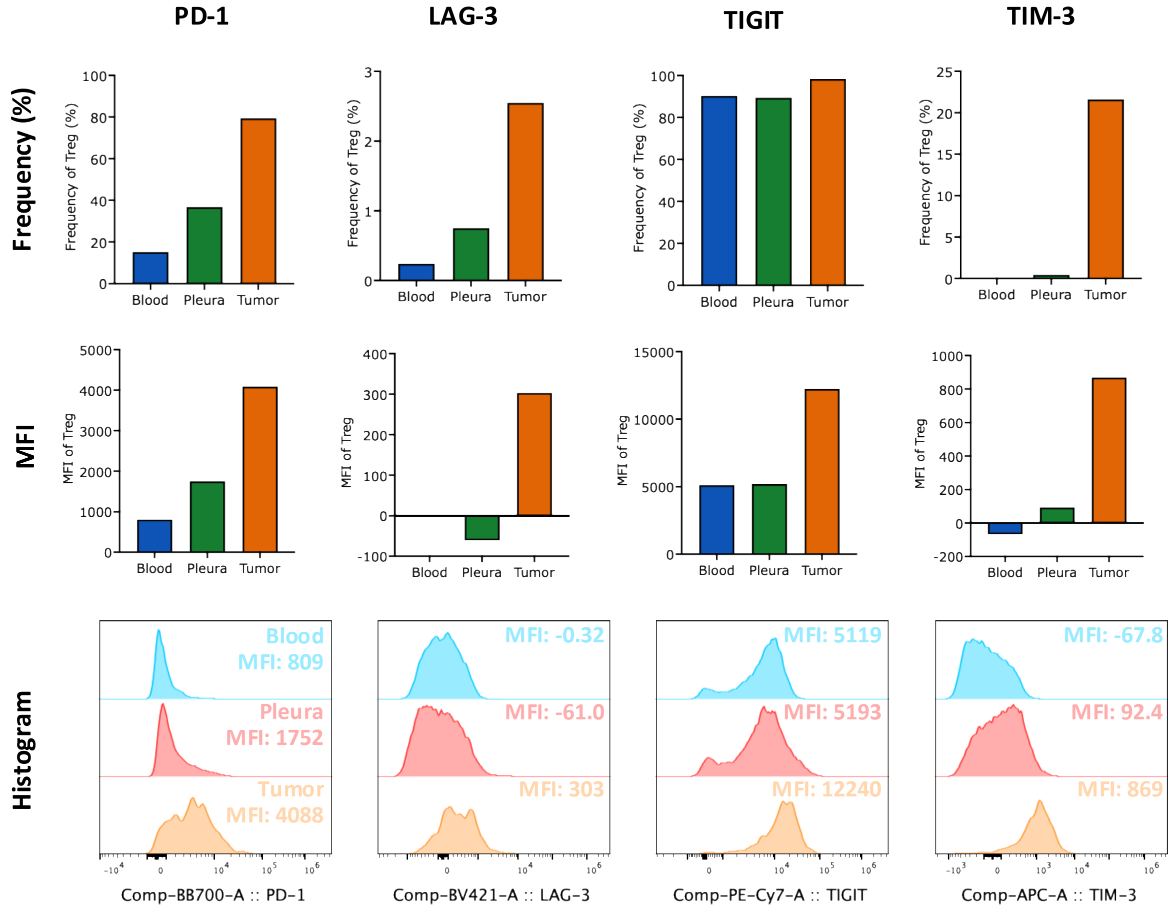


Frequency, mean fluorescence intensity (MFI), and histograms showing expression of *PD-1*, *LAG-3*, *TIGIT*, and *TIM-3* on regulatory $CD4^{+}CD25^{high}CD127^{low}$ (Treg) T cells across tissue compartments

### Figure S8

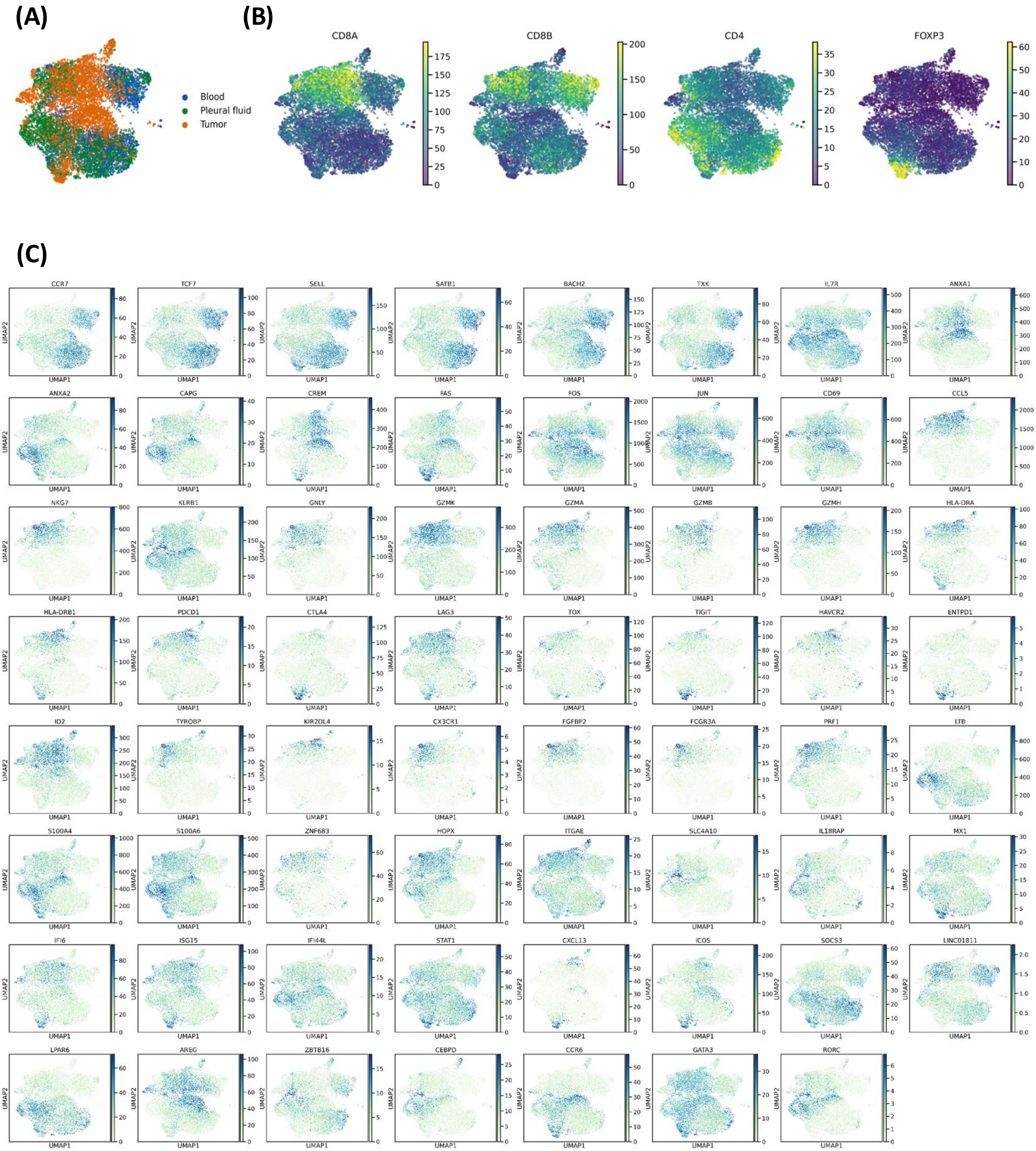


**(A)** UMAP visualization cells across MPE, blood and tumor **(B)** UMAPs showing expression of canonical T cell markers **(C)** UMAPs showing expression of curated markers

### Figure S9

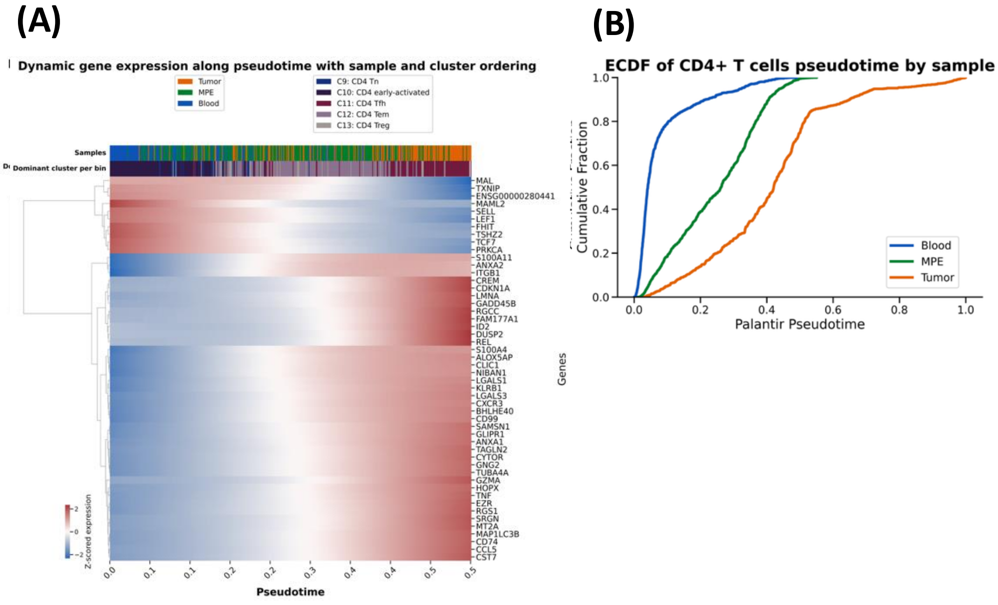


**(A)** Differential gene expression along pseudotime trajectory of CD4$^{+}$ T cells **(B)** Empirical cumulative distribution function (ECDF) of compartment-specific CD4$^{+}$ T cells along the pseudotime trajectory.

### Figure S10

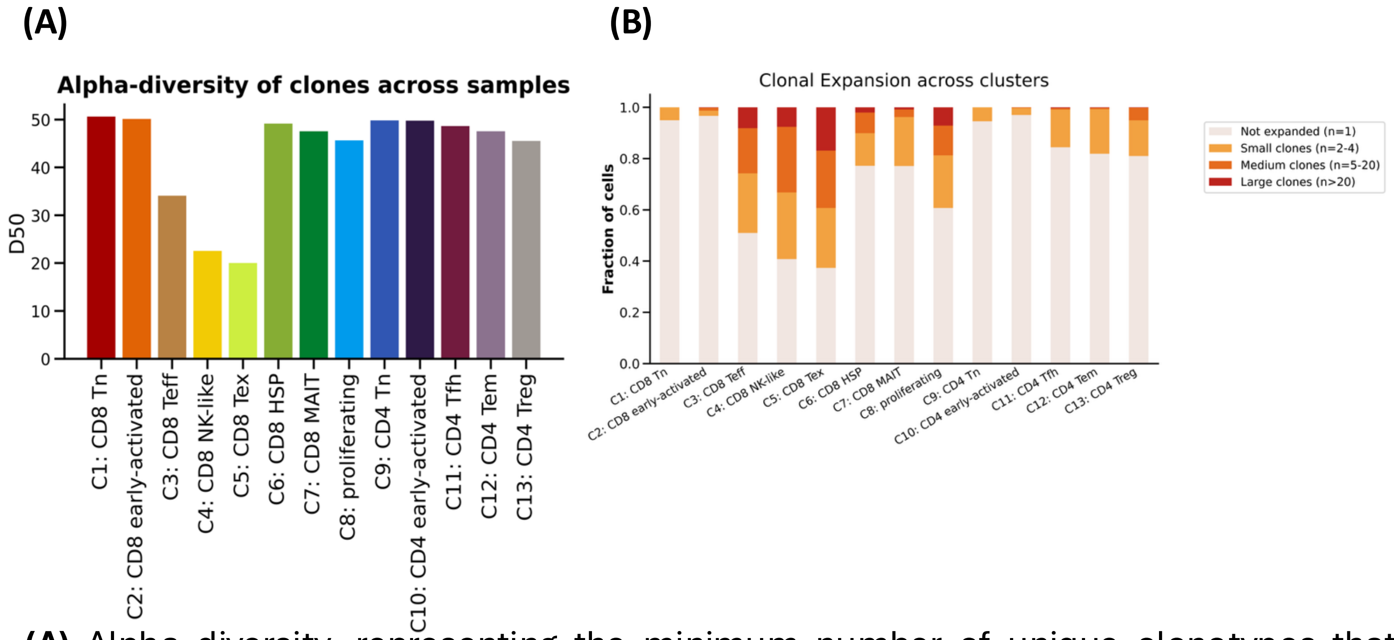


**(A)** Alpha diversity, representing the minimum number of unique clonotypes that account for 50% of all cells. across clusters. **(B)** Clonal expansion across Clusters.

### Figure S11

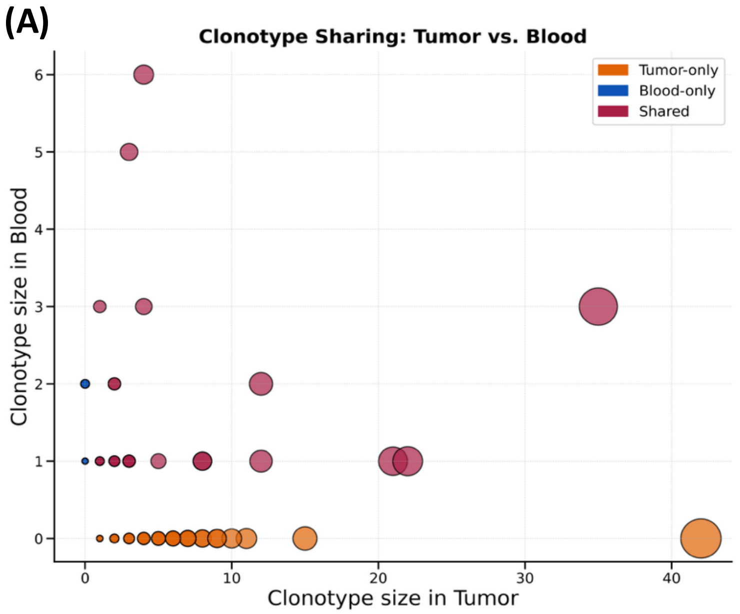


**(A)** Clonotype sharing between tumor and blood.

### Figure S12

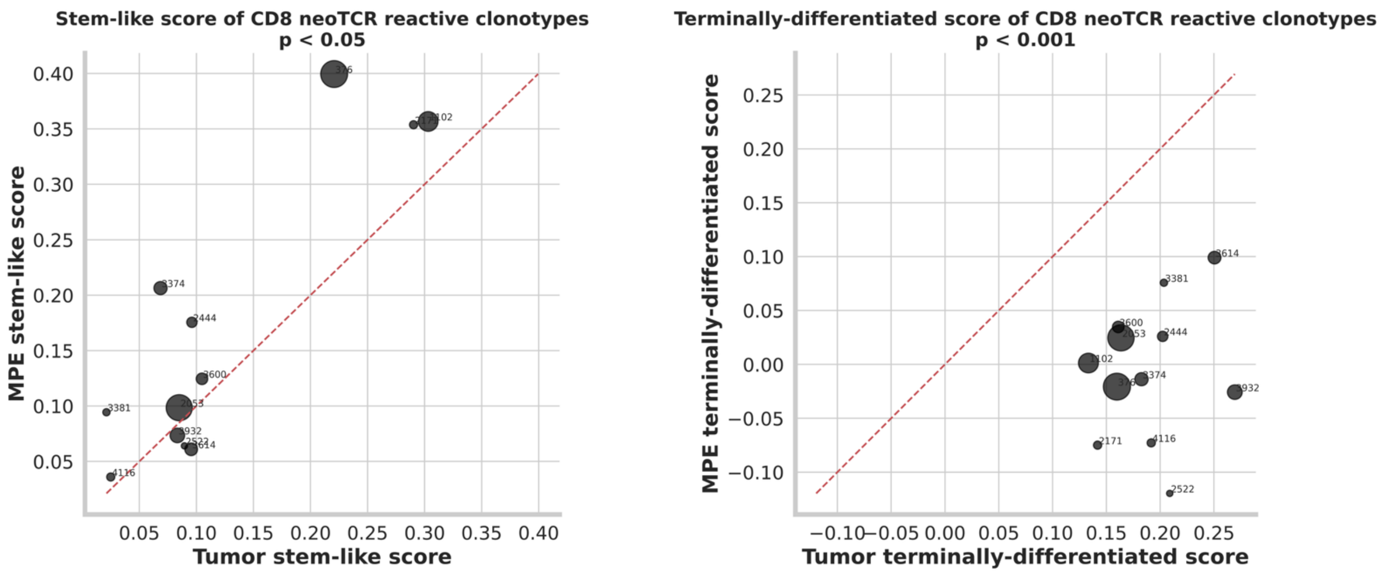


**(A)** Gene signature scores for stem like and **(B)** terminally differentiated T cells.

**Figure S13**

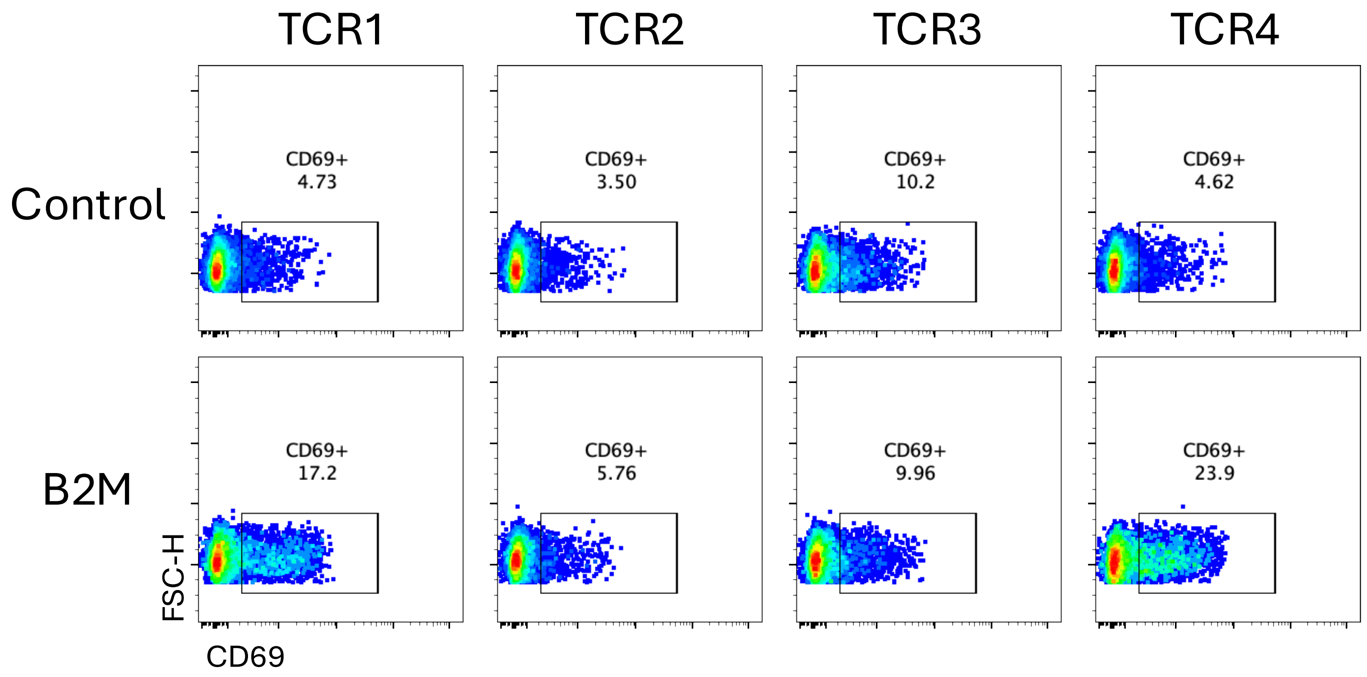


Representative flow cytometry plots of CD69 expression in Jurkat cells when co-cultured with cancer cell lines established from the patient's resected lung metastasis. The Jurkat cells are expressing TCR's identified in clonally expanded T cells identified in both tumor and MPE (TCR1–TCR4). The cell lines were either untransduced (top row) or transduced to re-express *B2M* (bottom row).

### Figure S14

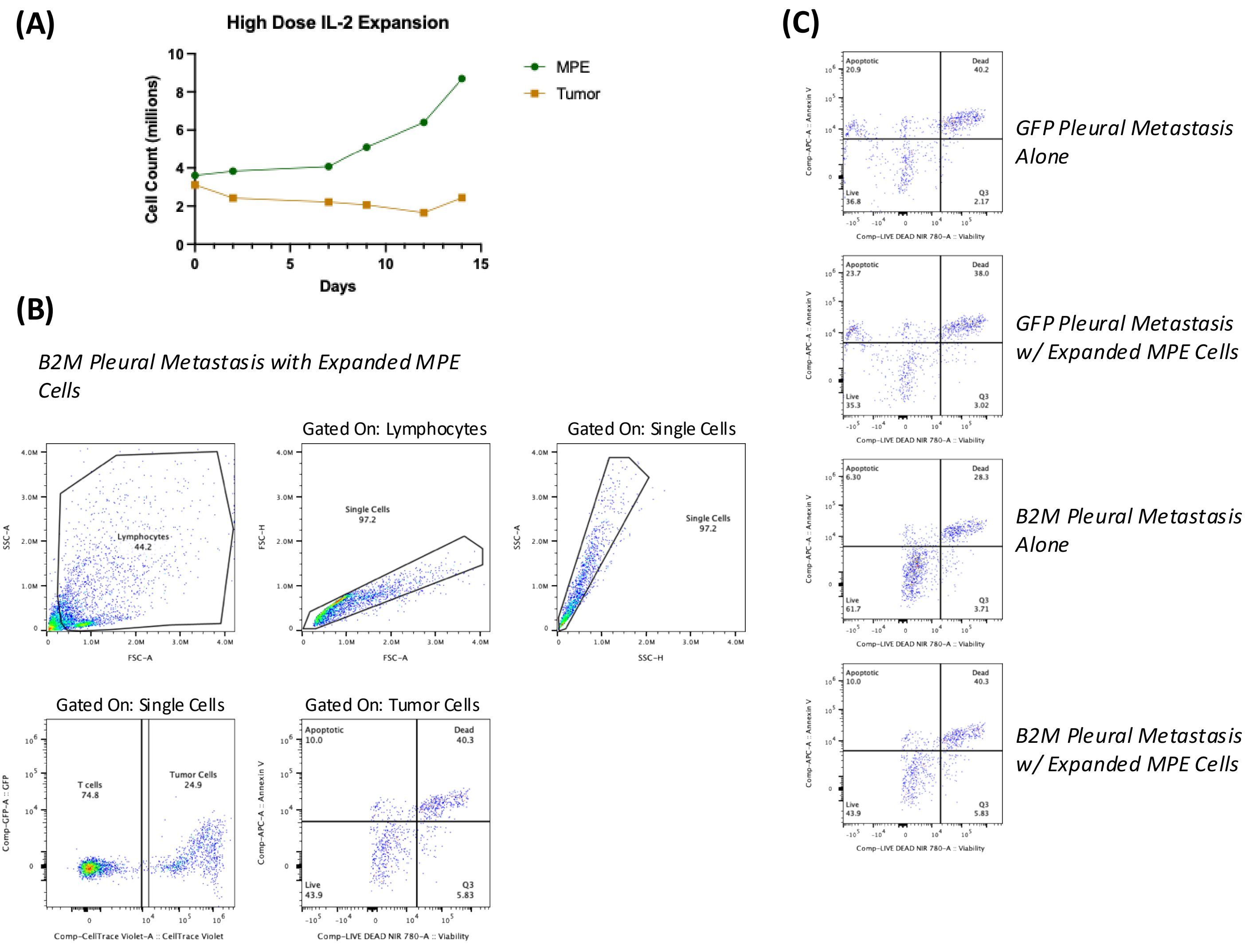


(A) Cell counts of MPE and tumor cells in high-dose IL-2 over the course of 2 weeks.
(B) Gating strategy for determining pleural metastasis cell viability in co-culture assay.
(C) Representative staining patterns of Live/Dead and Annexin V in GFP or B2M transduced pleural metastasis cells alone or with expanded MPE *CD8*$^{+}$ T cells
